# LncRNA-UCA1 Alleviates Septic Acute Kidney Injury through Regulating Endoplasmic Reticulum Stress in LPS-treated HK-2 Cells

**DOI:** 10.1101/2021.04.06.438568

**Authors:** TT Yu, FL Cai, J Niu

## Abstract

**Objective:** Septic acute kidney injury (AKI) is an important cause of death in patients with sepsis. This study sought to explore the function of the long noncoding RNA, urothelial carcinoma associated 1 (lncRNA-UCA1), in septic AKI and determine the underlying molecular mechanism.

**Methods:** HK-2 cells were treated with lipopolysaccharide (LPS) to establish an *in vitro* model of septic AKI. Quantitative reverse-transcription polymerase chain reaction (qRT-PCR) was used to detect the expression of lncRNA-UCA1. CCK-8 assay was used to detect the viability of HK-2 cells. Western blotting was utilized to examine protein expression. The contents of SOD, GSH, MDA, and ROS were determined using commercial kits. The apoptosis rate was calculated using TUNEL staining and flow cytometry.

**Results:** LncRNA-UCA1 was down-regulated in LPS-treated HK-2 cells. LPS significantly reduced the content of SOD and GSH in HK-2 cells, increased the production of MDA and ROS, and led to an increase in the rate of apoptosis. However, overexpression of lncRNA-UCA1 protected HK-2 cells from oxidative stress and apoptosis. Furthermore, LPS induced endoplasmic reticulum (ER) stress in HK-2 cells, which was inhibited by overexpression of lncRNA-UCA1.

**Conclusion:** Overexpression of lncRNA-UCA1 inhibited LPS-induced oxidative stress and apoptosis of HK-2 cells by suppressing ER stress.

## Introduction

Sepsis refers to systemic inflammatory response imbalance syndrome caused by severe infection. Inflammatory cytokines, cell growth factors, and other factors participate in sepsis, causing some patients to experience different organ dysfunctions and even septic shock symptoms[1]. Therefore, in recent years, sepsis has become widely recognized as a series of organ dysfunction diseases caused by severe inflammatory reactions from severe infections, resulting in disorders of the body’s state[2]. Although there have been great advances made in the treatment of infections, the incidence and mortality of sepsis patients have not declined significantly[3].

The most serious clinical complication of patients with sepsis is acute kidney injury (AKI). AKI occurs in 41.5% to 53.2% of patients with sepsis[4]. Among AKI patients admitted to intensive care units (ICU), sepsis is the most common cause, and AKI patients with sepsis account for 32.4% to 47.5% of all AKI patients[5]. In addition, the mortality rate of such patients is significantly increased, ranging from 50.6% to 70.2%, and the risk of later chronic renal disease is also significantly increased among survivors[6]. AKI in patients with sepsis results in a substantial economic burden to society and patients. Thus, there is an urgent need for new ways to treat septic AKI.

In the past, noncoding RNAs (ncRNAs) were considered “ junk”, but numerous studies have shown that such RNAs play important roles in biological processes[7]. Recently, increasing evidence has shown that ncRNAs are closely related to the progression and recovery of AKI. Some ncRNAs have also been suggested as diagnostic markers and therapeutic targets for AKI patients[8].

Long noncoding RNAs (LncRNAs) are ncRNAs longer than 200 nucleotides[9]. A large number of studies have shown that lncRNAs play important roles in various biological processes, such as epigenetic regulation, the cell cycle, development, and differentiation, which suggests that lncRNAs may also play roles in the progression of AKI[10]. Studies have shown that the upregulation of lncRNA-NEAT1 aggravated lipopolysaccharide (LPS)-induced AKI[11]. Additionally, Liu et al. showed that downregulation of lncRNA-TUG1 could promote the progression of septic AKI[12]. LncRNA-UCA1 has been studied extensively in tumors, and it is involved in multiple biological processes, such as tumor cell proliferation, apoptosis, and migration[13,14]. In addition, lncRNA-UCA1 can inhibit cardiomyocyte apoptosis, and thus, reduce myocardial ischemia-reperfusion injury. However, the role of lncRNA-UCA1 in septic AKI has not been studied.

In this study, we established an *in vitro* model of septic AKI. We revealed that the expression of lncRNA-UCA1 was decreased in septic AKI, and overexpression of lncRNA-UCA1 significantly inhibited LPS-induced oxidative stress and apoptosis of HK-2 cells. Thus, LncRNA-UCA1 may be a therapeutic target for septic AKI.

## Materials and methods

### Cell culture and transfection

HK-2 cells were purchased from the American Type Culture Collection (ATCC) and cultured in complete medium containing Dulbecco’s Modified Eagle’s Medium/F-12 (Gibco, Rockville, MD, USA), 10% fetal bovine serum (FBS) (Gibco, Rockville, MD, USA) and 1% penicillin / streptomycin (Gibco, Rockville, MD, USA). Cells were incubated at 37°C in 5% CO_2_ and 21% O_2_. To establish the *in vitro* model of septic AKI, HK-2 cells were treated with LPS (500 ng/mL) for 24 hours.

To upregulate the expression of lncRNA-UCA1 in HK-2 cells, adenoviruses (Ribobio, Guangzhou, China) carrying the full-length LncRNA-UCA1 gene sequence (AD-UCA1) or a GFP vector (Ad-GFP) were constructed. Ad-UCA1 or AD-GFP were transfected into HK-2 cells as directed by the manufacturer’s instructions.

### The qRT-PCR analysis

TRIzol reagent (Invitrogen, Carlsbad, CA, USA) was used to isolate the total RNA from HK-2 cells. The concentration and purity of the isolated RNA were detected using a NanoDrop One/Onec Spectrophotometer. The PrimeScript™RT reagent Kit (Takara, Japan) was used to synthesize lncRNA-UCA1 cDNA and the AceQ Universal SYBR qPCR Master Mix (Vazyme, Nanjing, China) was used to perform qRT-PCR. GAPDH was the internal control. The amplification primers of LncRNA-UCA1: forward primers(5’>3’): TTTGCCAGCCTCAGCTTAAT; reverse primers(5’>3’): TTGTCCCCATTTTCCATCAT. The amplification primers of GAPDH: forward primers (5’>3’): ACAACTTTGGTATCGTGGAAGG; reverse primers(5’>3’): GCCATCACGCCACAGTTTC.

### Superoxide Dismutase (SOD) Activity Assay

HK-2 cells were transfected with AD-UCA1 or AD-GFP and treated with LPS. The content of SOD in cells was determined using the SOD detection kit (Jiancheng Bioengineering Institute, Nanjing, China), according to the manufacturer’s protocols.

### Glutathione (GSH) Activity Assay

The content of GSH in HK-2 cells treated with LPS was detected using the GSH detection kit (Jiancheng Bioengineering Institute, Nanjing, China) according to the manufacturer’s protocols.

### Detection of Malondialdehyde (MDA) levels

The levels of MDA in HK-2 cells were examined using the MDA detection kit (Jiancheng Bioengineering Institute, Nanjing, China) according to the manufacturer’s protocols.

### Reactive Oxidative Stress (ROS) Activity Assay

The DHE-ROS kit (Bestbio, Shanghai, China) was used to detect ROS production in HK-2 cells. In cells, DHE is oxidized by ROS to ethidium oxide, which produces red fluorescence that was observed under an inverted fluorescent microscope. The fluorescence intensity was proportional to the level of ROS production.

### CCK-8 assay

We took log phase HK-2 cells and adjusted the concentration of cell suspension to about 5 × 10^4^/ml and add 100 μL into each well of a 96-well plate. After 24 hours, the supernatant in the well was aspirated, the cells were washed twice with PBS, and different concentrations of LPS (100, 200, 300, 400, 500, 600 ng/mL) was added to the wells for 24 hours. After that, the supernatant was discarded and the cells were wash twice with PBS. Next, the CCK-8 reagent (10 μL) (MCE, Nanjing, China) and serum-free medium (90 μL) were added for 1 hour. The absorbance at 450 nm was measured by a microplate reader.

### Flow Cytometry

HK-2 cells were treated as described above. The cells were collected after digestion using trypsin. We centrifuged the cell suspension for 5 minutes, then discarded the supernatant, washed with PBS, and centrifuged again in the same manner, repeating twice. Then, 100 μL of Binding Buffer was used to resuspend the cells. After that, 5 µL of Annexin V-FITc (KeyGen, Shanghai, China) and 5 µL of propidium iodide (PI) (KeyGen, Shanghai, China) was added in per tube. Finally, the apoptotic cells were detected using flow cytometry.

### Western blotting

Total protein in HK-2 cells was isolated using RIPA lysis buffer (Beyotime, Shanghai, China). The concentration of the isolated proteins was determined using the BCA method. Proteins were separated on 10% SDS-PAGE gels and transferred to polyvinylidene fluoride (PVDF) membranes. After the membranes were incubated in 5% skim milk, they were incubated with primary antibodies (pro-Caspase-3, Abcam, rabbit, 1:10,000; Caspase-3, Abcam, rabbit, 1:500; CHOP, Abcam, rabbit, 1:1,000; ATF4, Abcam, rabbit, 1:1,000; GRP78, Abcam, rabbit, 1:1,000; GAPDH, Abcam, rabbit, 1:1,000). Following incubation with primary antibodies, membranes were incubated with secondary antibodies for 1 h and visualized using a chemiluminescence system.

### TUNEL staining

HK-2 cells were fixed using 4% paraformaldehyde, treated with 0.1% Triton X-100 for 20 minutes, and stained using the One Step TUNEL Apoptosis Assay Kit (Beyotime, Shanghai, China) at 37°C for 1 hour, according to the manufacturer’s protocols. DAPI (Beyotime, Shanghai, China) was used to stain the nucleus and all staining was observed using a fluorescence microscope.

### Statistical analysis

Statistical analyses were performed using Statistical Product and Service Solutions (SPSS) 22. 0 software (IBM, Armonk, NY, USA). Data were represented as the mean ± standard deviation (SD). The *t*-test was used for analyzing measurement data. Differences between two groups were analyzed using the Student’s *t*-test. Comparisons between multiple groups were performed using one-way ANOVA, followed by post-hoc tests (Least Significant Difference). A p < 0.05 indicated significant differences.

## Results

### LncRNA-UCA1 was down-regulated in HK-2 cells treated with LPS

To construct a cell model of septic AKI, different concentrations (0, 100, 200, 300, 400, 500, 600 ng/mL) were used to treat HK-2 cells. And the viability of HK-2 cells was detected using CCK-8 assay. When the concentration of LPS was 500 ng/mL, cell viability decreased by about 50%, so the concentration was used in subsequent experiments (Fig. 1A). Then, we detected the expression of lncRNA-UCA1 using qRT-PCR and found that LncRNA-UCA1 was down-regulated in HK-2 cells treated with LPS (Fig. 1B). To study the role of lncRNA-UCA1, AD-UCA1 was transfected into the cells, and after which, the expression of lncRNA-UCA1 was significantly increased (Fig. 1C).

**Fig 1.**
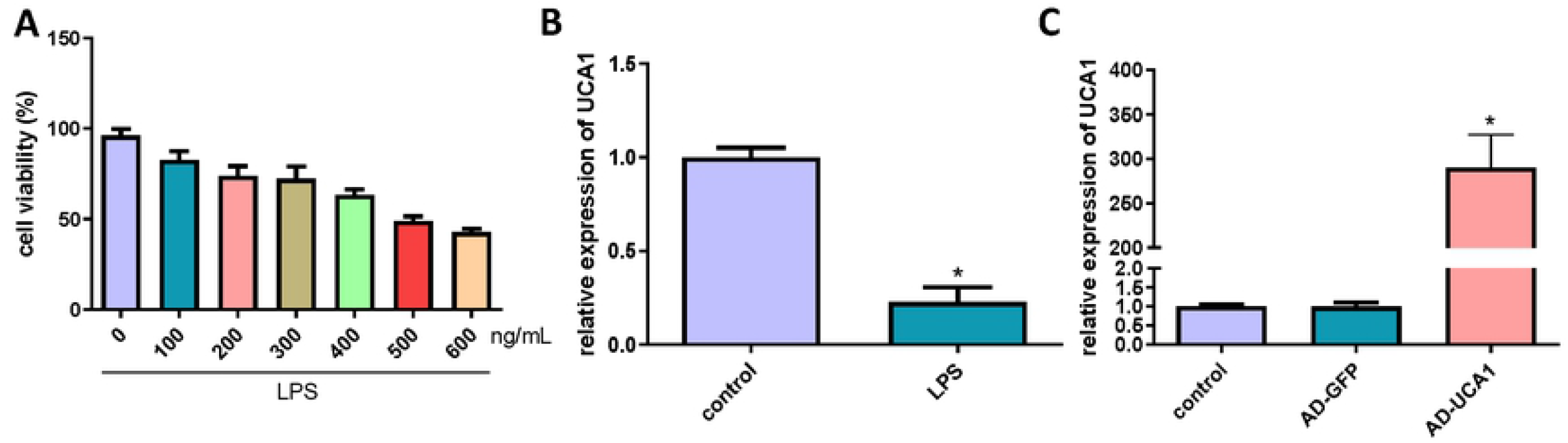
LncRNA-UCA1 was down-regulated in HK-2 cells treated with LPS. (A) The viability of HK-2 cells was detected using CCK-8 assay. (B) The expression of lncRNA-UCA1 in HK-2 cells treated with LPS was detected by qRT-PCR (“*” p < 0.05 vs control, n = 3). (C) Transfection of AD-UCA1 significantly increased lncRNA-UCA1 expression (“*” p < 0.05 vs AD-GFP, n = 3).

### LncRNA-UCA1 inhibited LPS-induced oxidative stress in HK-2 cells

We used commercial kits to detect the levels of antioxidant factors (SOD, GSH) in HK-2 cells. The treatment of LPS significantly reduced the content of SOD and GSH in HK-2 cells, while the overexpression of lncRNA-UCA1 markedly increased the levels of those factors (Fig. 2A and B). Moreover, LPS induced the production of MDA in cells, while overexpression of lncRNA-UCA1 remarkably inhibited its production (Fig. 2C). In addition, lncRNA-UCA1 significantly inhibited the production of ROS in cells induced by LPS (Fig. 2D and E). Thus, based on the above results, lncRNA-UCA1 attenuated the oxidative stress induced by LPS in HK-2 cells.

**Fig 2.**
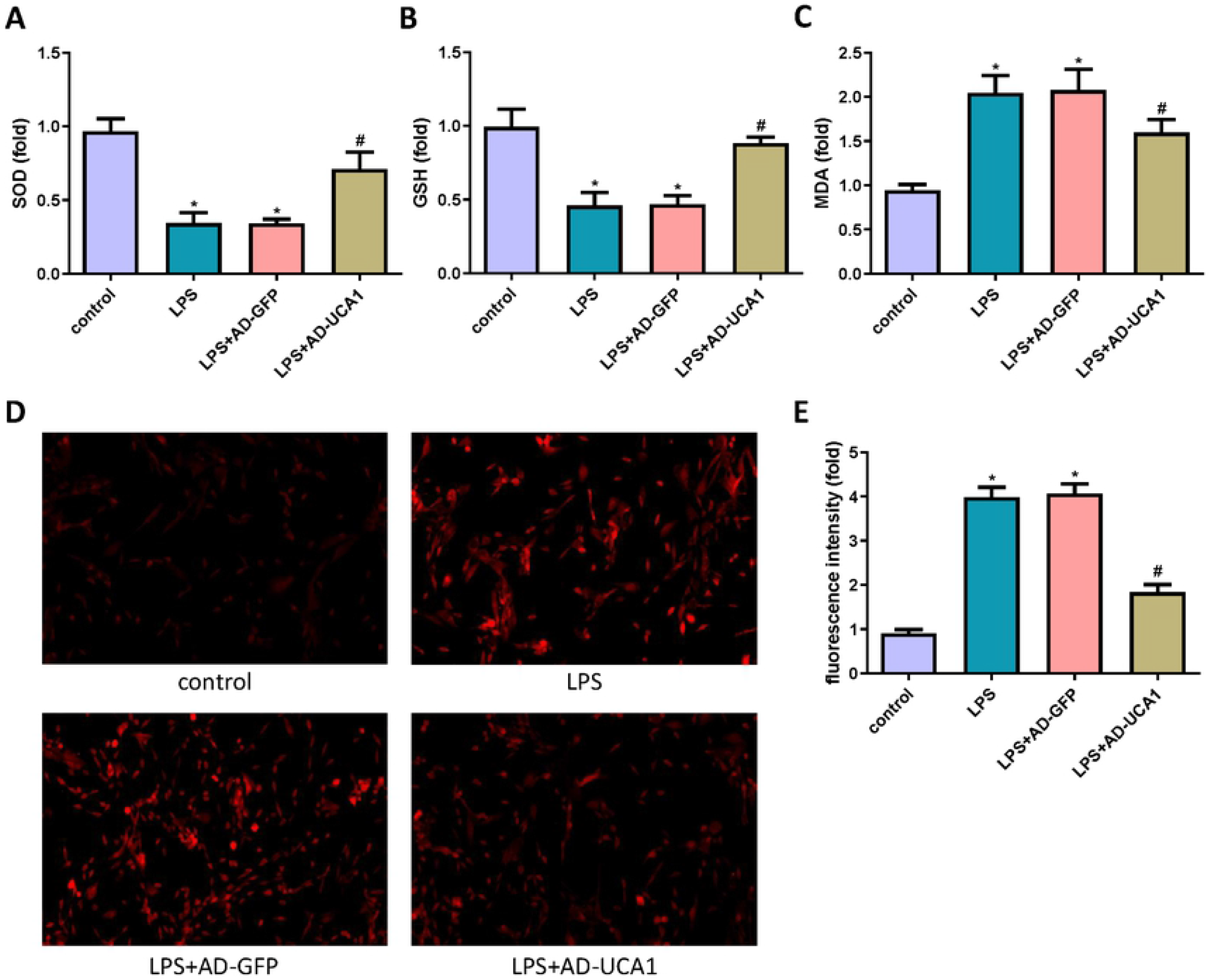
LncRNA-UCA1 inhibited LPS-induced oxidative stress in HK-2 cells. (A) The levels of SOD in HK-2 cells were detected (“*” p < 0.05 vs control, “#” p < 0.05 vs LPS+AD-GFP, n=3). (B) The levels of GSH in HK-2 cells were detected (“*” p < 0.05 vs control, “#” p < 0.05 vs LPS+AD-GFP, n=3). (C) The contents of MDA in HK-2 cells were detected (“*” p < 0.05 vs control, “#” p < 0.05 vs LPS+AD-GFP, n=3). (D) The DHE fluorescent probe was performed. (E) Statistical results of relative fluorescent intensity (“*” p < 0.05 vs control, “#”p < 0.05 vs LPS+AD-GFP, n=3).

### LncRNA-UCA1 inhibited LPS-induced apoptosis in HK-2 cells

The expression of the apoptosis-related protein, Caspase-3, and Kim-1 was detected by western blotting (Fig. 3A). Compared with the control group, the expression of Kim-1 in the LPS-treatment group was significantly increased. Compared with the LPS+AD-GFP group, the expression of Kim-1 in the LPS+AD-UCA1 group was significantly decreased (Fig. 3B). The change of Cleaved Caspase-3 expression was consistent with Kim-1 (Fig. 3C). In addition, overexpression of lncRNA-UCA1 can significantly reduce the decline of HK-2 cell viability induced by LPS (Fig. 3D). In addition, overexpression of lncRNA-UCA1 can significantly reduce the decline of HK-2 cell viability induced by LPS (Fig. 3E). Consistent with this, LPS significantly increased the rate of TUNEL positive cells, while overexpression of lncRNA-UCA1 significantly reduced this (Fig. 3F and G).

**Fig 3.**
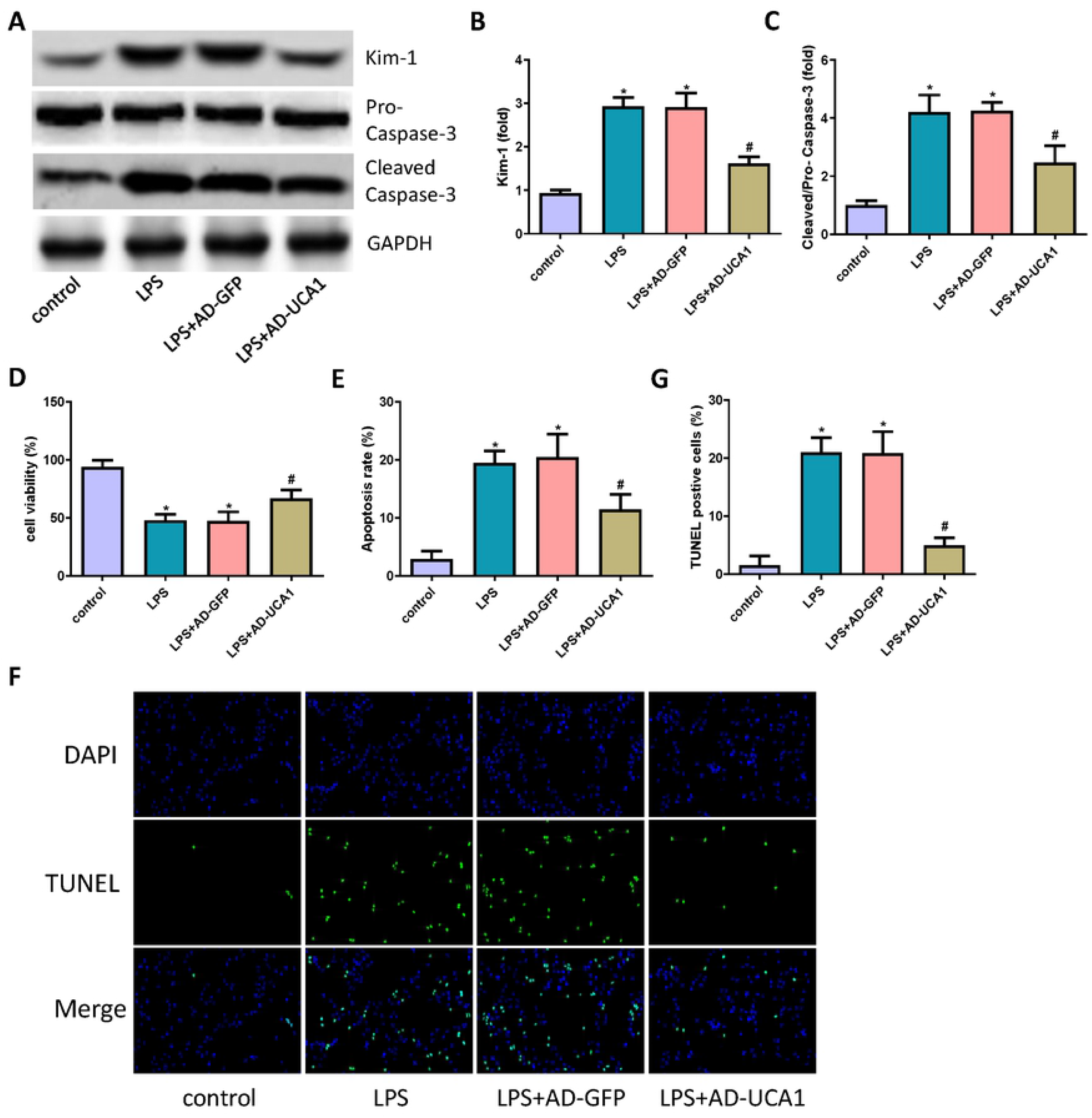
LncRNA-UCA1 inhibited LPS-induced apoptosis in HK-2 cells. (A) The expression of Cleaved Caspase-3 and Kim-1 in HK-2 cells were detected through western blot. (B and C) Statistical results of expression of Cleaved Caspase-3 and Kim-1 (“*” p < 0.05 vs control, “#” p < 0.05 vs LPS+AD-GFP, n=3). (D) The viability of HK-2 cells was detected (“*” p < 0.05 vs control, “#” p < 0.05 vs LPS+AD-GFP, n=3). (E) The apoptosis rate of HK-2 cells was detected using flow cytometry (“*” p < 0.05 vs control, “#” p < 0.05 vs LPS+AD-GFP, n=3). (F) Results of TUNEL staining of HK-2 cells (200×) (G) Statistical results of TUNEL staining (“*” p < 0.05 vs control, “#” p < 0.05 vs LPS+AD-GFP, n=3).

### LncRNA-UCA1 attenuated LPS-induced ER stress in HK-2 cells

To study whether lncRNA-UCA1 is involved in the mechanism of LPS-induced ER stress in HK-2 cells, ER stress biomarkers were detected through western blotting (Fig. 4A). Notably, LPS treatment increased the expression of CHOP, ATF4, and GRP78; however, lncRNA-UCA1 inhibited the expression of those proteins (Fig. 4B∼D). Thus, those results indicated that lncRNA-UCA1 protected HK-2 cells from LPS-induced ER stress.

**Fig 4.**
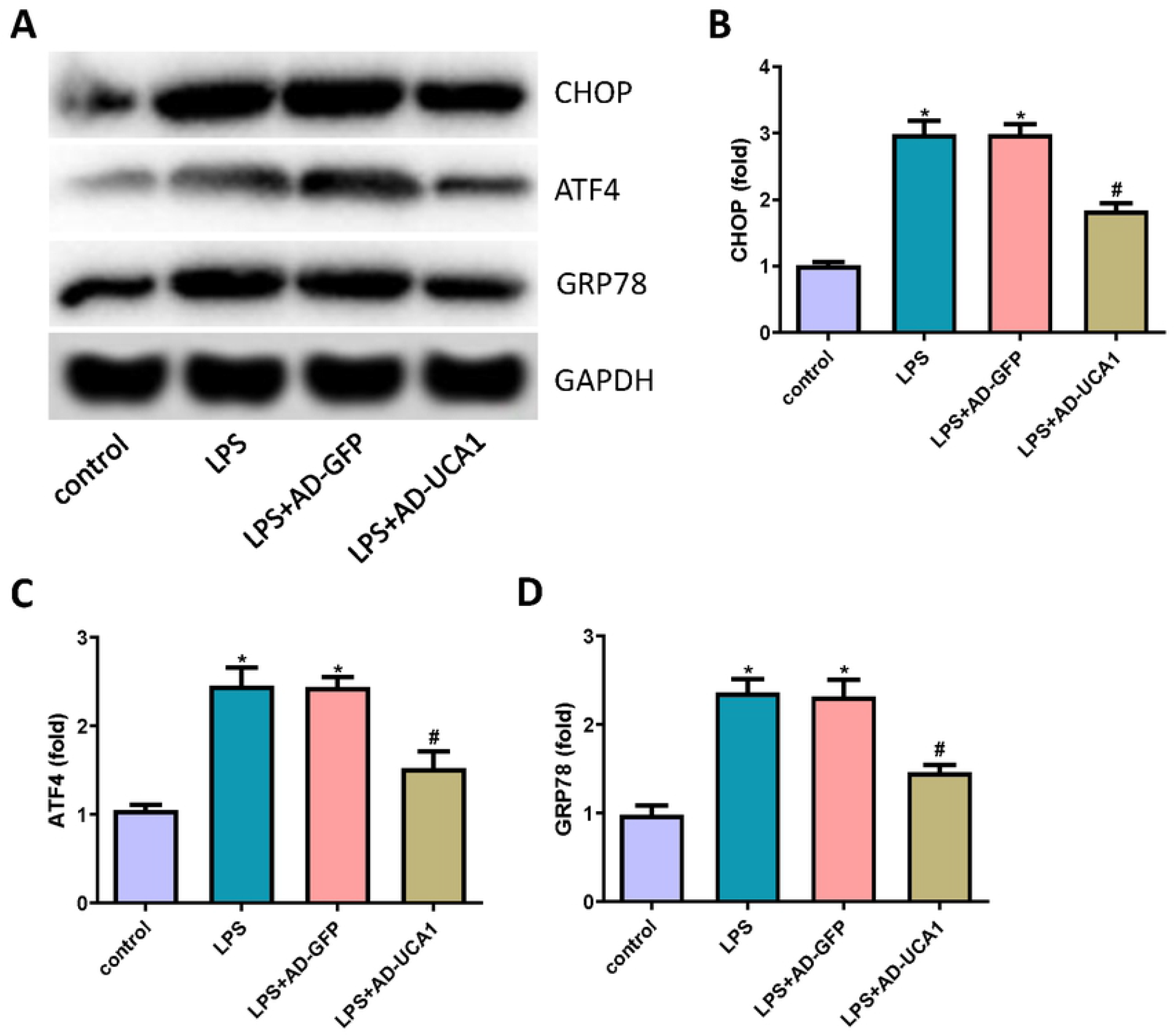
LncRNA-UCA1 attenuated LPS-induced ER stress in HK-2 cells. (A) The expression of CHOP, ATF4, and GRP78 in HK-2 cells were detected through western blot. (B∼D) Statistical results of expression of CHOP, ATF4, and GRP78 (“*” p < 0.05 vs control, “#” p < 0.05 vs LPS+AD-GFP, n=3).

## Discussion

In this study, we mainly revealed the following two points. First, the expression of lncRNA-UCA1 was decreased in the *in vitro* model of septic AKI. Second, overexpression of lncRNA-UCA1 inhibited oxidative stress and apoptosis of HK-2 cells by inhibiting the ER stress induced by LPS. The high mortality of septic AKI is due to the unclear pathogenesis of the disease. The pathogenesis of septic AKI is complex, with the involvement of a series of pathological processes, including renal hemodynamic disorders, necrosis and apoptosis of endothelial and epithelial cells, oxidative stress, and excessive immune and inflammatory reactions[15-17].

When septic AKI occurs, excessive inflammation induces the production of ROS. The ROS in the body attacks the polyvalent unsaturated fatty acids in the biofilm, thereby producing a large amount of MDA. The MDA level, thus, reflects the severity of the ROS attack on the body’s cells[18]. SOD is a specific enzyme responsible for the removal of superoxide free radicals. GSH is an important member of the non-enzymatic antioxidant system, which removes oxygen free radicals from cells. Collectively, the contents of SOD and GSH have become important indicators of the body’s antioxidant capacity.

Apoptosis refers to programmed death, which can be triggered via the intrinsic pathways, mitochondrial pathway, and ER pathway. Current research shows that the apoptosis of renal tubular epithelial cells plays an important role in septic AKI[19,20]. In addition, multiple drugs are also being studied to inhibit Caspase cascades, reduce the apoptosis of renal tubular epithelial cells, and play a role in protecting the kidney[21]. These further confirmed the dominant role of renal tubular epithelial cell apoptosis in the mechanism of septic AKI.

In this study, LPS was used to establish a septic AKI model of HK-2 cells. After LPS treatment of HK-2 cells, the content of MDA in cells increased, while SOD activity and GSH expression decreased. Those findings suggested that the production of ROS in septic AKI was increased, the endogenous enzymatic and non-enzymatic antioxidant effects were weakened, and strong oxidative stress appeared. However, the overexpression of lncRNA-UCA1 inhibited the above changes. In addition, LPS induced the production of cleaved Caspase-3 in HK-2 cells, thereby inducing cell apoptosis. Again, the overexpression of lncRNA-UCA1 significantly inhibited the apoptosis of HK-2 cells.

Under pathological conditions, multiple stimulating factors can cause unfolded or misfolded protein aggregation in the ER, which causes ER dysfunction and triggers ER stress. Excessive ER stress activates the transcription of the pro-apoptotic coding gene, CHOP, thereby activating Caspase-12 and inducing cell apoptosis. In this study, LPS induced the expression of ER stress markers, which indicated the occurrence of ER stress. In addition, CHOP was expressed in large amounts, thereby inducing apoptosis. Similarly, the overexpression of lncRNA-UCA1 significantly inhibited those processes.

This study had some limitations. First, the mechanism by which lncRNA-UCA1 functions has not been well studied. At present, lncRNA is mainly thought to regulate biological functions by adsorbing microRNAs (miRNA), however, the present study did not investigate the target miRNAs of lncRNA-UCA1. Second, this study lacked *in vivo* experiments to evaluate the protective effect of lncRNA-UCA1 on septic AKI.

## Conclusions

In summary, lncRNA-UCA1 was down-regulated in septic AKI and the overexpression of lncRNA-UCA1 inhibited LPS-induced oxidative stress and apoptosis in HK-2 cells by suppressing ER stress.

## Acknowledgements

Not applicable

## Availability of data and materials

The datasets used and/or analyzed during the current study are available from the corresponding author on reasonable request.

## Authors’ contributions

FC and TY designed the study. TY and JN performed the experiments. TY analyzed the data. FC and TY wrote the manuscript.

## Abbreviations

AKI: acute kidney injury
UCA1: urothelial carcinoma associated 1
LPS: lipopolysaccharide
qRT-PCR: quantitative reverse-transcription polymerase chain reaction
ER: endoplasmic reticulum
FBS: fetal bovine serum
AD-UCA1: adenoviruses carrying the full-length LncRNA-UCA1 gene sequence
Ad-GFP: adenoviruses carrying a GFP vector
SOD: Superoxide Dismutase
GSH: Glutathione
MDA: Malondialdehyde
ROS: Reactive Oxidative Stress
CCK-8: cell counting kit-8
TUNEL: terminal deoxynucleotidyl transferase (TdT)-mediated dUTP nick end labeling.

